# The quasi systematic nature of splitter cells

**DOI:** 10.1101/2024.06.07.597927

**Authors:** Naomi Chaix-Eichel, Snigdha Dagar, Fréderic Alexandre, Thomas Boraud, Nicolas P. Rougier

**Author notes:** These authors contributed equally to this work (Naomi Chaix-Eichel, Snigdha Dagar). These authors also contributed equally to this work (Fréderic Alexandre, Thomas Boraud, Nicolas P. Rougier).

## Abstract

During the past decades, hippocampal formation has undergone extensive studies, leading researchers to identify a vast collection of cells with functional properties. Several investigations, supported by carefully crafted models, have examined the origin of such cells. The most recent models hypothesize that temporal sequences underlie the observed spatial properties. We aim at investigating whether a random recurrent structure is sufficient to allow such latent sequence to appear. To do so, we simulated an agent with egocentric sensory inputs that must navigate and alternate choices at intersections. We were subsequently able to identify several splitter cells inside the model. Remarkably, when we systematically lesioned the identified splitter cells, the model’s behavioral performance remained intact: in the vast majority of cases, new splitter cells re-emerged through network reorganization, while in the remaining cases, the task was solved without any detectable splitter cells, demonstrating that splitter cells are not necessary to the task resolution. Position, orientation, and decision representations could also be successfully decoded from the reservoir activity, even after repeated lesioning. Subspace alignment analysis further revealed that this reorganization preserves the task-relevant population geometry while redistributing activity within the null sub-space, with the trajectory-encoding dimension rotating in neuron space across lesions. Together, these findings demonstrate that splitter cell activity is primarily task-driven and does not derive from a specific architecture or learning rule: splitter cells emerge generically across random recurrent networks that successfully solve the task, across a broad and robust range of dynamical parameters, and are not necessary for task performance. Our results therefore challenge the notion of functional necessity for specific neural populations.

## 1 Introduction

The hippocampal formation has been heavily studied in the past few decades (Maguire et al., 1996; Buzsáki and Moser, 2013; Hartley et al., 2014; Moser et al., 2017) and re-searchers have since then established a huge repertoire of cells displaying very specific properties. In his book “The Brain from Inside out” (György Buzsáki, 2019), György Buzsáki named a few of them in a footnote (page 356): *place cells, time cells, grid cells, head direction cells, boundary vector cells, border cells, object cells, object vector cells, distance cells, reward cells, concept cells, view cells, speed cells, memory cells, goal cells, goal direction cells, splitter cells, prospective cells, retrospective cells, island cells, ocean cells, band cells, object cells, pitch cells, trial order cells, etc*. Each and every cells have been characterized in terms of correlation between their activity and some combination of high level property involving space, time and internal state. This is the case for place (cell fires when animal is in a specific place), head direction (cell fires preferentially when head is pointing toward a specific direction), time cells (cell fires at successive moments), etc. Facing such huge repertoire, one may legitimately wonder how the brain orchestrates all these information and makes use of it to ensure survival. Part of the answer is given by György Buzsáki himself in the same footnote where he further explained that *physiological attributes of neurons in the hippocampal-enthorinal system … might be explained by the apparent distinctiveness of a few sets of continuous distributions*. In other words, even though we can and we do observe these cells *in vivo*, they might be a simple epiphenomenon: their activity might be correlated to some unknown latent variables.

This is actually hardly different from the hypothesis provided by Raju et al. (2024) where authors suggest that spatial representations are not explicitly encoded but emerge as a byproduct of sequence learning. Their model is based on a variant of Hidden Markov Model (HMM) theory and employs a clone-structured causal graph (CSCG) (George et al., 2021) to differentiate between various sequences of input. In this framework, multiple “clones” represent the same observation across different contexts, enabling the model to handle ambiguous sensory input sequentially. Consequently, the representation of space arises implicitly through the process of sequence learning, rather than being an explicit function of the hippocampus. Other research has similarly challenged the traditional space-centric view of the hippocampus. Using the concept of successor representation (Dayan, 1993), Stachenfeld et al. (2017) suggest that hippocampal cells encode a predictive map rather than a purely euclidean spatial map. Sanders et al. (2020) propose that the hippocampus builds more abstract representations of environmental structure based on the animal’s subjective beliefs rather than directly representing objective spatial properties of the environment. Whittington et al. (2020) introduced the Tolman-Eichenbaum machine (TEM), a model that unifies spatial and non-spatial functions of the hippocampus. They argue that hippocampal cells emerge as a result of the system learning to represent abstract relationships, enabling the representation of both spatial and non-spatial information. Luo et al. (2024) demonstrated that units resembling place, head-direction, and border cells arise spontaneously in deep neural networks trained purely on a non-spatial task (object recognition), and even in entirely untrained networks with random weights, suggesting that spatial cell types are an inevitable emergent property of any sufficiently complex information processing system rather than a specialization unique to the hippocampal formation. In this study, we align with current approaches and propose to reconsider the role of hippocampal cells. More precisely, we decided to focus our study on splitter cells, which encode information about past or future trajectories (Wood et al., 2000). In other words, they demonstrate different firing patterns depending on the animal’s origin or intended destination, even if the animal is in the same physical location, as explained in Figure 1.

**Figure 1:**
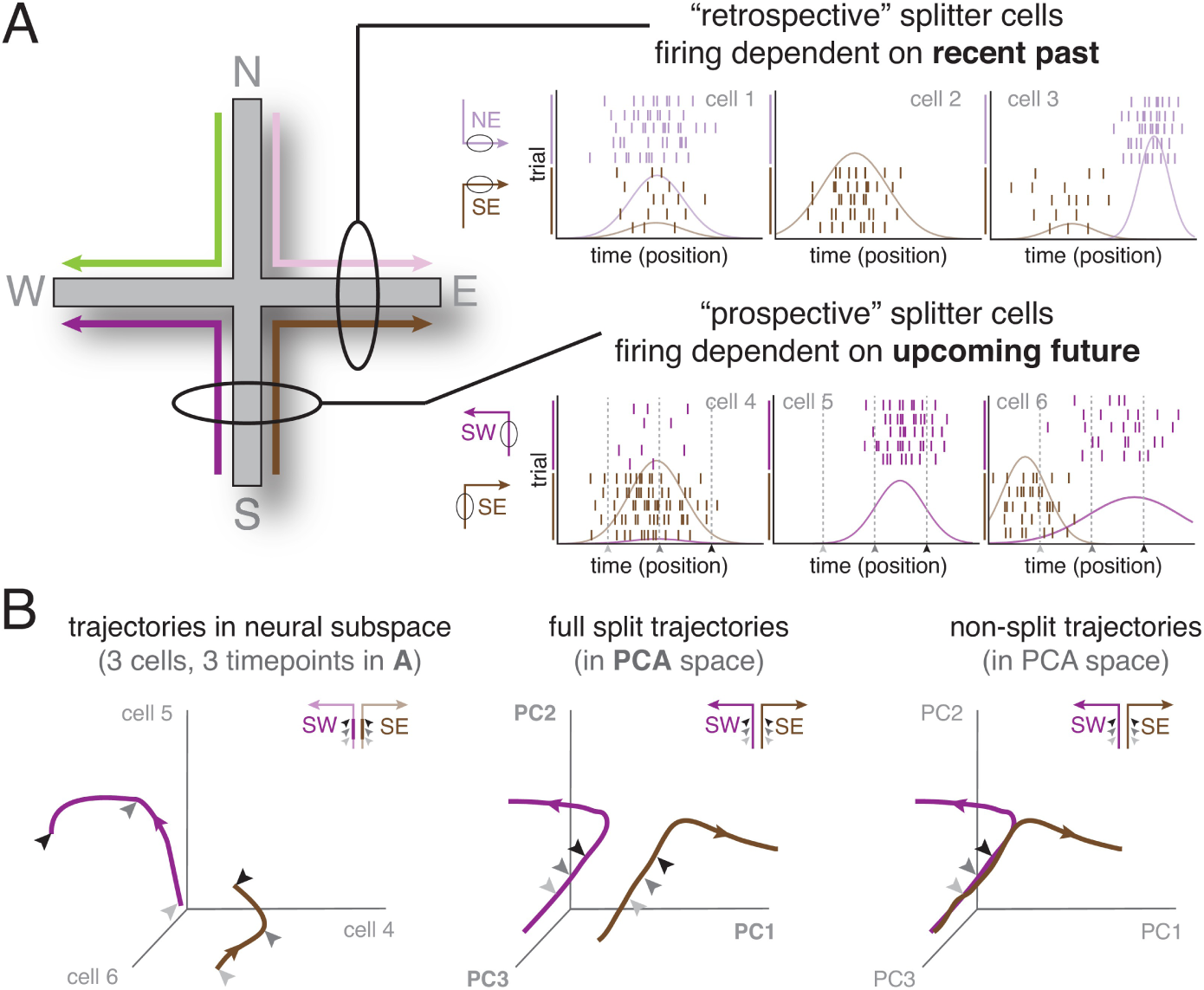
Figure from Duvelle et al. (2023) explaining splitter cells at the single cell and population levels. (A) Schematic activity of six idealized splitter cells during a plus-maze task with four possible trajectories (SW, SE, NW, NE). Splitter cells differentiate either past or future trajectories. Some show rate differences (left), others respond to only one trajectory (middle), or shift their firing location depending on trajectory (right). (B) Left: Schematic neural trajectories in population activity space for SW and SE paths. Despite identical physical paths, activity diverges based on future direction (E vs W). Middle: Neural trajectories projected onto the first three principal components, showing separation during both shared and diverging path segments. Right: Hypothetical population activity without splitter cell modulation, shown for comparison.

We adopt a novel perspective based on György Buzsáki’s interpretation of the brain as a large reservoir of neurons with pre-existing patterns of activity (György Buzsáki, 2019). According to this view, when a new experience occurs, the brain assigns it to a pre-existing pattern from a vast reservoir of internally generated patterns, thereby assigning behavioral meaning to the neuronal sequence. This mechanism is considered more biologically plausible than creating entirely new patterns for each experience. Building on this concept, we propose to study a reservoir computing model composed of a pool of recurrently interconnected neurons, representing a vast reservoir capable of generating an extensive repertoire of neuronal patterns. The model’s synaptic weights are randomly initialized and kept fixed, implying that the neuronal patterns within the reservoir remain unchanged after learning. The only trained part is the readout layer that connects the reservoir to the output. This fixed-weight approach does not model plasticity in the reservoir, but we argue that it is sufficient and consistent with György Buzsáki (2019)’s perspective. This is supported by experiments showing that learning only modestly and transiently impacts network dynamics (Golub et al., 2018). Furthermore, this view of the navigation system allows for a simple model that is easy to understand and manipulate, which is our main novelty and contribution compared to the other models listed: we don’t require complex architectures or concepts to study splitter cells, just a pool of randomly interconnected neurons. Despite its simplicity, this model has already demonstrated great success and robustness in several works by Eric A. Antonelo (Antonelo et al., 2007, 2008; Antonelo and Schrauwen, 2009, 2012), and has even been successfully implemented on real robots navigating spatial environments (Antonelo and Schrauwen, 2015). We demonstrate that this model can robustly solve a T-maze alternation task, specifically within a continuous state-space. Similarly to the model of Raju et al. (2024), our model handles ambiguous sensory inputs reflecting a more realistic scenario. Rather than relying on allocentric information such as spatial coordinates, the model has access only to partially observable inputs, i.e. egocentric information: distance sensors. These sensors can read identical values in different locations within the maze, making the sensory information ambiguous. This feature increases the task’s realism and difficulty and distinguishes our approach from Stachenfeld et al. (2017); Whittington et al. (2020); George et al. (2021), where such ambiguity was not explicitly addressed.

By conducting a theoretical analysis, we show that the internal states of certain model units exhibit properties similar to biological splitter cells. Interestingly, when we ‘lesioned’ the model by explicitely removing these units, the performances remained unaffected and new splitter cells emerged. In a configuration where navigation behavior is forced, this lesion–regeneration cycle can be repeated multiple times, leading to a reorganization of the internal dynamics of the network, that is shown to be able to support the task. This repeated emergence suggests that splitter cell-like representations are not fixed structural features but functional strategies that arise from the demands of navigation. Although it remains unclear whether splitter cells drive navigation or emerge as a consequence of it, our findings caution against attributing functional necessity to any specific unit subset. Instead, the network flexibly reallocates computation across remaining or newly initialized units. Furthermore, position, orientation and decision can be successfully decoded from the model, even after lesions, aligning with prior studies (Luo et al., 2024) but using a more parsimonious architecture consistent with György Buzsáki (2019). The flexibility of splitter cell emergence indicates that these representations are likely byproducts of the navigation task, rather than innates structural modules dedicated to trajectory coding.

## 2 Materials and Methods

### 2.1 Environment

#### Task presentation

The class of tasks called spatial alternation has been widely used to study hippocampal and working memory functions (Frank et al., 2000). For the purpose of our investigation, we simulated a continuous versions of the task, wherein an agent must navigate through an 8-shaped track and alternate between right and left choices at a decision point, then returns to the central corridor, essentially following an 8-shape trace while moving (see the ‘LR’ task of Figure 2-A). This 8-maze environment offers an ideal setup for observing splitter cells when the agent enters the central corridor, because the agent can have various past trajectories leading to the central corridor and different future trajectories upon exiting it. A more challenging variant requires the agent to alternate every two loops (see ‘LL-RR’ task in Figure 2-B), generating left-left-right-right sequences. This variation allows trajectories with the same past but different futures (e.g., right-left vs. right-right) or different pasts but the same future (e.g., right-right vs. left-right), allowing to distinguish between retrospective splitter cells (encoding past trajectories) and prospective splitter cells (encoding future trajectories).

**Figure 2:**
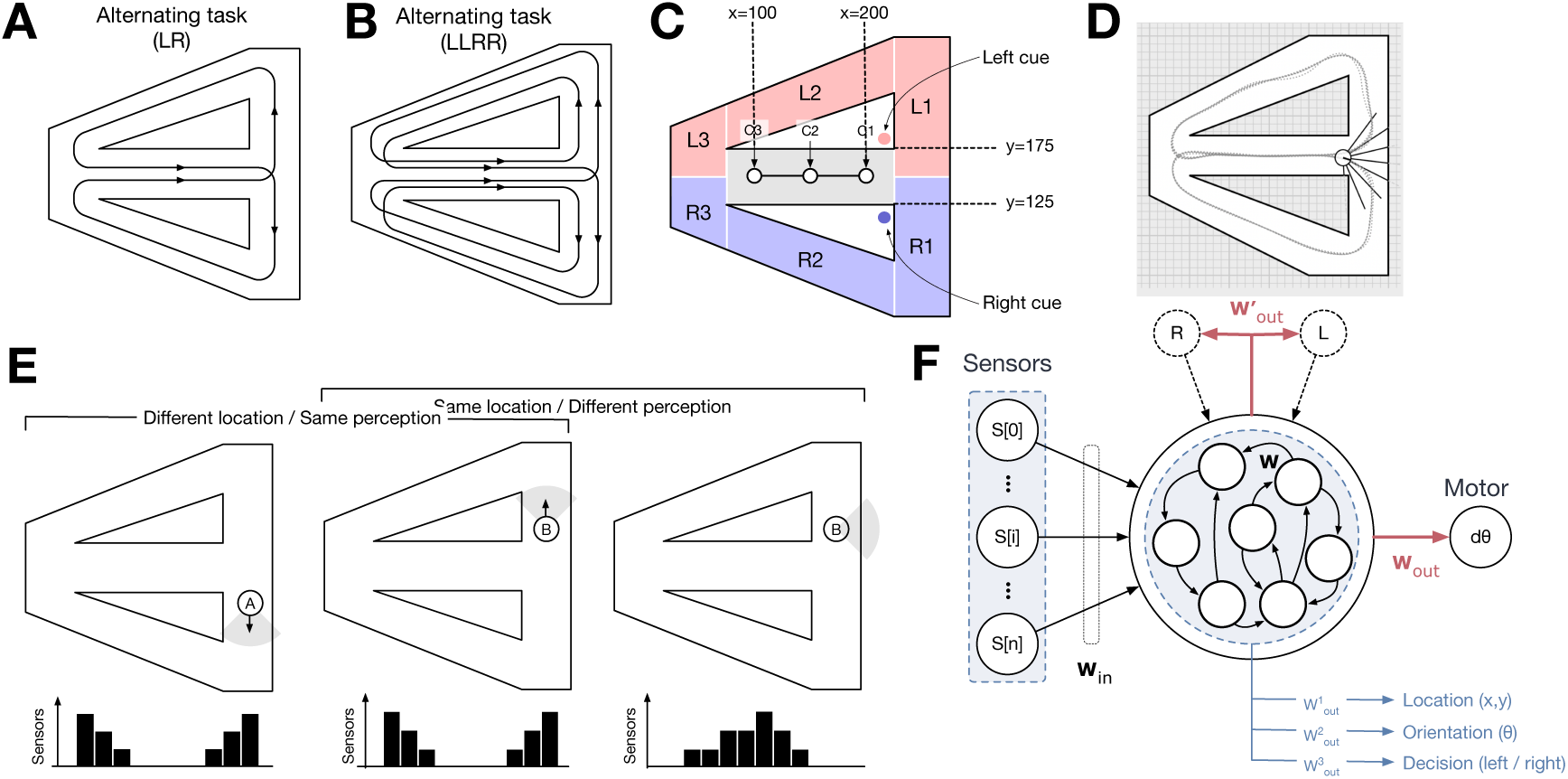
**A-B.** An expanded view of a T-Maze. **A**. The first task involves alternating every time it is located at the decision point, consequently doing the sequence of decision ‘R-L-R-L-R…’. **B**. The second task involves alternating every two times, consequently doing the sequence of decision ‘R-R-L-L-R-R-…’. **C. Recording area of the splitter cells.** In the red zone, the agent commits to left (L); in the blue zone, to right (R). The grey region in the central corridor indicates the area where splitter cell activity is recorded. Markers C1, C2, and C3 correspond to specific locations used to track neural activity within this recording zone. The left and right cues act as salient signals presented when the bot enters the central corridor, indicating which direction to take at the upcoming decision point. Unlike externally provided cues, these signals are generated internally by the model itself. **D. Snapshot of simulation.** Each ray represents a distance sensor reading from the bot, and the dotted line indicates its trajectory. These visualizations allow to assess whether the model successfully completes the task. **E. Ambiguous egocentric sensory inputs during the agent simulation**. In several locations within the maze, such as when the bot is at positions A and B, the sensors record identical distance values despite the bot being in different positions. Additionally, there are instances where the bot, despite being at the same location, receives varying inputs, as seen in the comparison between positions B and C, illustrating the model’s challenge in handling ambiguous sensory information. **F. Model Architecture.** The model consists of 8 sensor inputs, a reservoir, and a motor output representing the change in relative orientation. Black arrows indicate fixed connections, while red arrows represent plastic, trainable connections. Two versions of the model: the *regular model* takes sensor inputs and outputs the next orientation change; and the *self-generated cues model* includes two readouts: one for predicting the next orientation change and another one for generating the next cue. This cue is then fed back into the model as an additional input. The model receives both sensor values and the self-generated cue as input.

### 2.2 Modeling framework

#### Reservoir computing

Solving an alternating task necessitates the presence of a functional working memory, as the agent must retain information about its previous direction, such as turning right in the prior loop, to inform its decision to turn left at the subsequent decision point. Reservoir computing is thus a natural choice because of its fading memory property Pascanu and Jaeger (2011). This enables the system to retain past experiences, facilitating their recall for subsequent decisions. Moreover, the reservoir computing approach has been shown to exhibit some biologically plausible characteristics (Lukoševičius and Jaeger, 2009). Consequently, our model consists of a reservoir computing network of type Echo State Network (ESN) (Jaeger, 2007) that controls the movement of the agent solving a continuous navigation task in the 8-maze of Figure 2-A/B. An ESN is a recurrent neural network (called reservoir) composed of randomly connected units, associated with an input and an output layer. As shown in Figure 2-F, inputs are projected into the reservoir, a nonlinear, high-dimensional space, enabling the integration of information across time and space. Only the output units, referred to as readout neurons (highlighted in red), are trained. The reservoir neurons follow the dynamics:

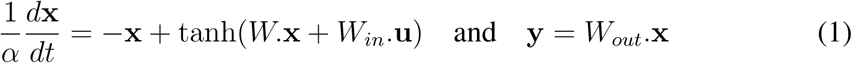

where **x**, **u** and **y** represent the reservoir states, input, and output. *W*, *W_in_*, and *W_out_* are weight matrices, while *tanh* refers to the hyperbolic tangent function. *α* refers to the leak rate, a crucial parameter of the ESN that plays a central role in controlling the memory and timescale of the network’s dynamics: a low leak rate indicates a longer memory and a slower dynamics, whereas a high leak rate leads to a shorter memory but a higher speed of update dynamics (Lukoševičius, 2012). The model was built thanks to the python library ReservoirPy (Trouvain et al., 2020).

#### Training

Only the output weights *W_out_* are trained, using supervised learning with linear ridge regression method (the Tikhonov regularization) on pre-generated data:

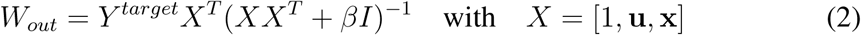

where *Y ^target^*, *β* and *I* are respectively the target signal to approximate, the regularization coefficient and the identity matrix.

To generate training data, we implemented a simple Braintenberg vehicle (Braitenberg, 1986) where the agent moves automatically with a constant speed and changes its orientation according to the values of its sensors. The model takes as input the sensor values *S*_0_*, …, S_n_* of the agent and outputs the next variation of orientation *dθ* (see Figure 2-F). At each time step the sensors measure the distance to the walls and the bot turns in order to avoid the walls. Since the Braitenberg algorithm is designed solely for obstacle avoidance, the desired trajectory is initially shaped by temporarily adding walls at key intersection points. These walls are then removed, and the agent repeats the trajectory without them, ensuring the correct sensory inputs are recorded in the absence of the guiding structures. The sensors are limited in their ability to calculate distance, saturating for distances exceeding 100 distance units. Additionally, the sensor readings are affected by Gaussian noise, represented as *N* (0, 0.03). A reading of 0 indicates proximity to an object, while a reading of 1 signifies that the object is far away or not detected. The sensors can record identical distance values even when the bot is not at the same spatial location in the maze, as illustrated in Figure 2-E. The challenge for the model lies in disambiguating this sensory information to accurately determine the bot’s location. The decision to use distance sensors as input provides a simplified egocentric representation of the environment, abstracting the complexities of the multiple sensory modalities involved in visual and auditory processing that animals like rats typically utilize. We argue that this simplification is justified: despite their simplicity, distance sensors still yield ambiguous sensory signals, preserving a key challenge of real-world navigation. Moreover, certain animals, such as bats (Wenstrup and Portfors, 2011) and others (Brinkløv et al., 2013; Evans, 1973), use echolocation to obtain distance-based information analogous to these sensor readings. Notably, egocentric boundary cells that likely contributing to hippocampal input in the medial entorhinal cortex (mEC), also encode similar spatial information (Long et al., 2025).

At each timestep, the position of the bot is updated as follows:

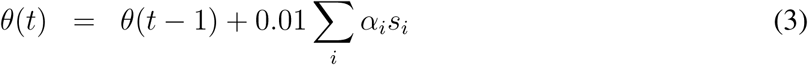

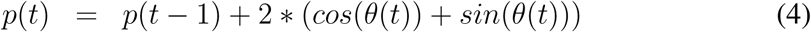

where *p*(*t*) and *p*(*t* + 1) are the positions of the agent at time step *t* and *t* + 1, *θ*(*t*) is the orientation of the agent, calculated as the weighted (*α_i_*) sum of the values of the sensors *s_i_*. The norm of the movement is kept constant and fixed at 2.

### The regular and the self-generated cue model

We introduce two model variants depicted in Figure 2-F. The regular model relies solely on sensory input from the bot to make decisions. Its variant, the self-generated cue model, includes two readouts: one for predicting the next change in orientation, and another for generating a cue. These cues, interpretable as visual indicators within the central corridor (see Figure 2-C), signal to the agent which direction to take at the upcoming decision point: “L” for left and “R” for right. In more classic setups, such cues are typically provided by the environment; here, they are self-generated by the model. They correspond to binary values, set to 1 when the agent is expected to turn in the corresponding direction, and are only active when the agent enters the central corridor. This self-generated cue is fed back into the system as an additional input, allowing the model to receive both the sensory data and the generated cue.

### 2.3 Analysis

#### Task solving

After the training phase, we look at the model’s ability to solve the task by directly looking at the online simulation whose snapshot is depicted in Figure 2-D. If the bot can perform the navigation task for at least five loops (one loop is composed of 700 time steps with one alternation, going one time to the right and one time to the left), the model is considered to have successfully solved the task.

#### Single-cell level analysis

We analyze the mean firing activity of individual neurons within the reservoir to identify splitter cells. These neurons exhibit different firing rates along overlapping segments of distinct trajectories. To detect such cells, we applied the ANCOVA (Analysis of Covariance) method described in Box 3 of Duvelle et al. (2023), originally used in hippocampal studies (Wood et al., 2000; Kinsky et al., 2020). This statistical technique tests for significant differences in firing rates across behavioral conditions. Neuronal firing are recorded as the agent traversed the central stem of the T-maze, a shared segment of both trajectories (see Figure 2-C). Firing rates are grouped by trajectory direction: right-to-left (RL), left-to-right (LR), right-to-right (RR) and left-to-left (LL). For each trajectory, we record at 15 activity sequences. The main ANCOVA factor was trajectory direction (x-axis in Figure 2-C), with the agent’s lateral position (y-axis) and orientation included as covariates. These covariates account for variability in position and heading along the central corridor. Speed was not included as a covariate, as it remained constant throughout the experiment. We classified neurons as splitter cells if the ANCOVA revealed a statistically significant difference in firing rates between RL and LR conditions with *p <* 0.0001. Detected splitter cells were visualized using raster plots of their firing activity recorded along the central corridor. Their activity was converted to dynamic firing rates using a Poisson process model (Heeger et al., 2000; Dayan and Abbott, 2005), with each neuron’s activity scaled accordingly:

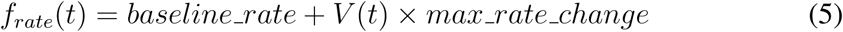

where *f_rate_*(*t*) corresponds to the computed firing rate at time *t*, *V* (*t*) is the normalized activity of the reservoir neuron at time *t*, *baseline rate* is the baseline firing rate (in spikes per second) and set to 45 and *max rate change* is the maximum deviation from the baseline also set to 45. The firing rate is subsequently converted into spike counts using a Poisson process:

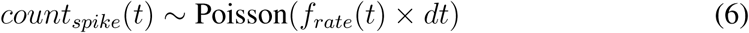

where *count_spike_*(*t*) represents the number of spikes occurring at time *t*, given the firing rate *f_rate_*(*t*). The Poisson process models the probabilistic occurrence of spikes over time with its paramater *λ* = *f_rate_*(*t*)*dt*.

### Population-level analysis

We analyze the reservoir state of the ESN at the population level using Principal Component Analysis (PCA) as a dimensionality reduction technique to identify patterns and significant features within the processed data.

#### Lesioning splitter cells within the model

We investigated the effects of lesioning the detected splitter cells within the model. To do so, we inactivate the connections associated with the splitter cells by setting their weight values to 0 in *W_in_*, *W*, and *W_bias_*. The resulting model is denoted as the *Lesioned model* (see Figure 6-D). We extended this analysis cumulatively: starting from the intact reservoir, we identified and lesioned all splitter cells, retrained the readout, and repeated this lesion-retrain cycle iteratively until task performance could no longer be recovered.

#### Subspace alignment analysis

To assess whether cumulative splitter-cell lesions perturb the full population geometry and whether splitter reemergence reflects preservation of task-relevant structure, we applied Singular Vector Canonical Correlation Analysis (SVCCA) (Raghu et al., 2017) to compare reservoir activity at baseline (iteration 0, no lesion) with each post-lesion iteration k. This method is well suited to populations whose effective size changes across lesions, as similarity is computed in a reduced activity space rather than in the full neuron basis. For each seed and lesion iteration, we extracted one reservoir activity vector per corridor crossing (mean firing over approximately 100 time steps in the central corridor zone), separately for LR and RL trajectories. Baseline and post-lesion activity matrices were compared on matched crossing conditions.

SVCCA proceeds in two steps. First, activations are mean-centered and Singular Value Decomposition (SVD) is applied to each representation independently: For each activity matrix **X**^LR^, the SVD reads:

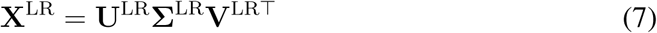

where **U**^LR^ contains the left singular vectors, **Σ**^LR^ is a diagonal matrix of singular values, and **V**^LR^ contains the right singular vectors. Retaining the top *K* left singular vectors *U^LR^_K_* ∈ *R^N×K^* yields the reduced representation:

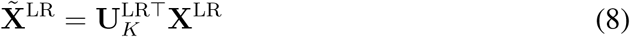

which expresses activity in SVD coordinates rather than neuron coordinates. The same reduction is applied to **X**^RL^, yielding **X**^° RL^.

Second, Canonical Correlation Analysis (CCA) (Hotelling, 1992) is applied and finds weight vectors *w^cca^_LR_, w^cca^_RL_* ∈ *R^N×K^* such as to maximize the following correlation across crossings:

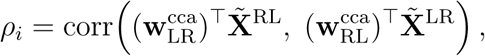

The CCA weights *w^cca^_LR_* are expressed in SVD coordinates, and the corresponding direction in neuron space is recovered by:

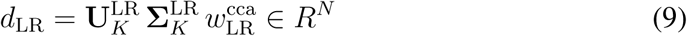

The task axis **d** ∈ *R^N^* is defined as:

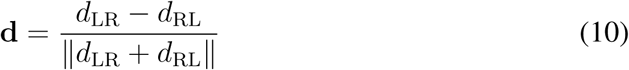

We additionally quantified the task-axis rotation relative to baseline by measuring |*cosθ*| between the task axis at baseline and at iteration k (Björck and Golub, 1973):

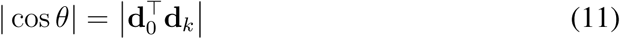

where **d**_0_ and **d***_k_* are unit vectors. A value of | cos *θ*| ≈ 1 indicates that the task axis is preserved in neuron space across the lesion; a value near 0 indicates that the task axis has rotated to a nearly orthogonal direction, meaning that different neurons contribute to trajectory encoding after the lesion.

### Decoding spatial information associated with other hippocampal cell types

In addition to splitter cells, we explore whether the reservoir’s dynamical systems could encode other hippocampal cell-related activities, such as head-direction, decision, and position. We employ an additional readout, referred as the “decoder readout”, trained using offline supervised learning to decode their activity. The decoder was trained on 7500 time steps and tested on 2500 time steps, with a regularization factor of 1*e* − 3 for all readouts. Head-direction cells, which fire in response to a specific orientation, are decoded by generating output vectors representing eight distinct orientations intervals: [(−*π,* −3*π/*4), (−3*π/*4, −*π/*2),(−*π/*2, −*π/*4), (−*π/*4, 0), (0*, π/*4), (*π/*4*, π/*2), (*π/*2, 3*π/*4), (3*π/*4*, π*)]. For example, if the bot’s orientation is between 3*π/*4 and *π*, the output vector will be [0, 0, 0, 0, 0, 0, 0, 1]. For decision, we generate training data by recording the bot’s next decision (right or left) at one of the intersection point following the central corridor, labeling these decisions as 0 (left) or 1 (right). For position, the training data was the sequence of coordinates of the bot’s trajectory. To complement this analysis, we tested whether spatial information could be directly decoded from raw sensory inputs to predict the robot’s position, orientation, or decision. This served as a control condition: if spatial variables can be decoded from sensory inputs alone, then successful decoding from the reservoir cannot be attributed to the reservoir’s computational properties per se, but may instead reflect information already present in the input stream.

## 3 Results

### 3.1 Model performances

We conducted four experiments in total: (1) the regular model on the ‘LR’ task, (2) the regular model on the ‘LLRR’ task, (3) the self-generated cue model on the ‘LR’ task, and (4) the self-generated cue model on the ‘LLRR’ task (see Figure 2). Each experiment was run with 10 different random seeds, corresponding to 10 distinct random initializations of the input (*W*_in_) and recurrent (*W*) connectivity matrices within the reservoir.

For the ‘LR’ task, both the regular and self-generated cue models successfully completed the task across all 10 seeds. Hyperparameters (detailed in the Tables section) were manually optimized and showed strong robustness, consistent with findings from Antonelo and Schrauwen (2015). Although we used 1000 reservoir units, similar performance was achieved with 500 units when other parameters were adjusted accordingly. Input/output connectivity could vary widely without affecting performance. The most sensitive parameters were the leak rate and the spectral radius, which jointly control the dynamics of the model, the memory, and the stability. Regularization was also important to prevent overfitting. For efficiency, the remainder of the analysis was conducted using the offline learning rule due to the slower convergence of online learning.

In the ‘LLRR’ task, the self-generated cue model succeeded consistently across all 10 seeds, whereas the regular model failed since it could navigate for only two loops (in average and below the five-loop success threshold). It is to be noted that this task is more demanding, as it requires remembering the last two decisions instead of just one, increasing the working memory demand. The self-generated cue model overcomes this limitation thanks to the presence of the internal cues, which act constantly to tell which decision to take next, and reduce the need for persistent memory within the reservoir.

In contrast, the regular model appears to lack sufficient working memory to solve the task.

### 3.2 Splitter cells

### 3.3 Splitting effect at the single cell level

We replicate the analysis from Duvelle et al. (2023) to assess whether their hypotheses about splitter cells also apply to our reservoir-based models. Splitter cells were identified in the regular model and, notably, also emerged in the self-generated cue model for both tasks. This is an interesting result, as the self-generated cue model does not rely on splitter cells to encode past or future directions: the cues explicitly inform the model at each time step in the central corridor which direction to take next, greatly reducing the need for working memory, and thus for cells encoding past and future trajectories. Yet, splitter cells still emerge, suggesting that their presence is not strictly tied to task demands. The average number of splitter cells for each model and task is shown in Table 2.

**Table 1:**
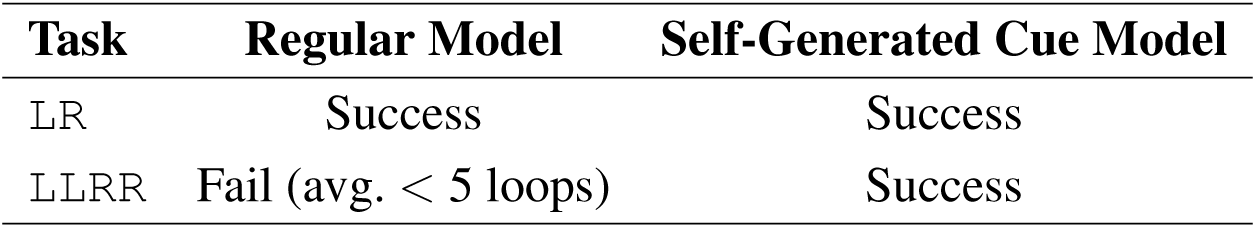
Task performance summary for each model. A task is considered successful if the model completes at least 5 loops on average across 10 seeds.

| **Task** | **Regular Model** | **Self-Generated Cue Model** |
| --- | --- | --- |
| LR | Success | Success |
| LLRR | Fail (avg. < 5 loops) | Success |

**Table 2:** Average number of splitter cells across 10 seeds for each model-task combination.

| **Task** | **Regular Model** | **Self-Generated Cue Model** |
| --- | --- | --- |
| LR | $217.3 \pm 148.2$ | $326.7 \pm 138.5$ |
| LLRR | — | $402.5 \pm 159.1$ |

Examples of splitter cell activity are shown in Figure 3-A, which displays the firing patterns of six splitter cells from the self-generated cue model as the bot traverses the central corridor (grey zone in Figure 2-B) during the LR task. Each trace shows the firing activity of a single splitter cell over 50 time steps within the corridor, averaged across 15 traversals per trajectory. Clear differences in activity can be observed: red traces correspond to trials where the bot moves from right to left (’RL’), and blue traces to left-to-right (’LR’) movements (see color code in Figure 2-B). Raster plots generated using a Poisson process, shown in the lower panels of Figure 3-A, further illustrate the average firing activity of this subset of splitter cells during central corridor traversal. Comparable patterns are observed with the regular model on the same task.

**Figure 3:**
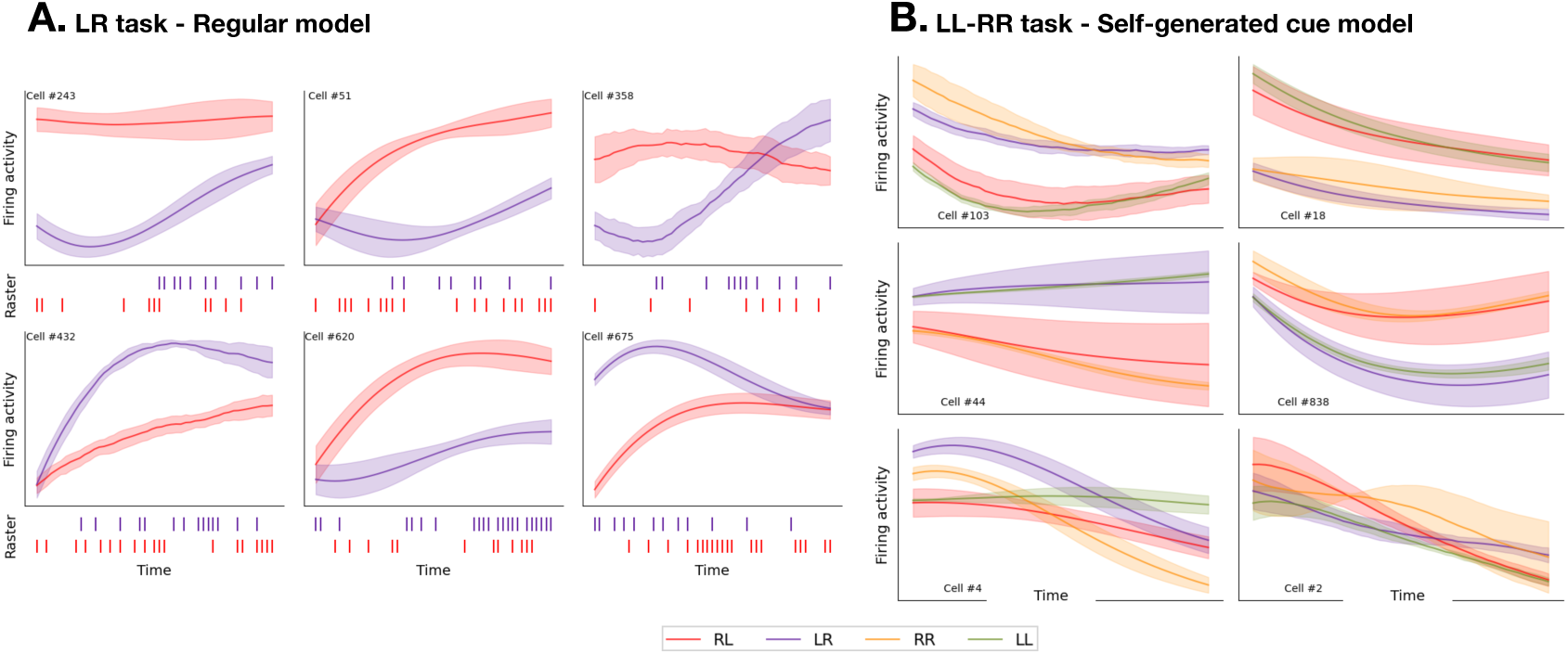
Comparing firing rates of individual splitter cells in the central stem. The identification of these cells was achieved using the ANCOVA method, which revealed significantly different firing activities (*p <* 0.001) in overlapping trajectories ‘RL’ (blue), ‘LR’ (red), ‘RR’ (yellow), ‘LL’ (green). The activity of splitter cells is recorded when the bot enters the central corridor (grey zone in Figure 2-B). Each trace represents the firing activity of a splitter cell in the central corridor over 50 time steps. Each line corresponds to the mean firing activity of at least 5 traversals, and the shaded areas indicates the standard deviation. Bellow each graph, the corresponding raster plots are depicted showing a difference in spiking activity rate between the overlapping trajectories during one traversal. **A**. Splitter cells in the regular model for the LR task. Comparable patterns are observed with the self-generated cue model on the same task. **B**. Prospective cells, retrospective cells and non splitter cell in the self-generated-cue model for the ‘LL-RR’ task. The two cells in the first row are prospective: they differentiate between trajectories that share the same past but have different futures (e.g., LR and RR vs. LL and RL). The two cells in the second row are retrospective: they differentiate between trajectories with the same future but different pasts (e.g., LR and LL vs. RL and RR). The last row shows two cells are not splitter cells, as they exhibit no detectable splitting effect across different trajectories. These patterns were not observed in the regular model on the same task, as the model failed to successfully complete it.

Figure 3-B shows similar results for the ‘RR-LL’ task, with the two first rows that shows the activity of four splitter cells from the self-generated cue model. Additionally, this task allows one to distinguish between prospective and retrospective splitter cells. Notably, the cells in the first row are prospective: they differentiate between trajectories that share the same past but have different futures (e.g., ‘LR’ and ‘RR’ (right-to-right) vs. ‘LL’ (left-to-left) and ‘RL’). In contrast, the cells in the second row are retrospective: they differentiate between trajectories with the same future but different pasts (e.g., ‘LR’ and ‘LL’ vs. ‘RL’ and ‘RR’). For comparison, the last row shows cells that are not splitter cells, as they exhibit no detectable splitting effect across different trajectories.

### 3.4 Splitting effect at the population level

#### Temporal Context Model versus Latent State Inference Model

The splitting effect refers to distinct neural trajectories in neural state space when the animal occupies the same spatial location. The hypothesized neural activity patterns during an alternation task are illustrated in Figure 1-B, where two distinct trajectories are shown: one corresponding to movement from the central corridor to the left arm, and the other from the same central corridor to the right arm. In the middle panel, these trajectories occupy separate regions in PCA space even though the animal is at the same spatial location, indicating a splitting effect. In contrast, the right panel illustrates the absence of a splitter signal: the neural trajectories fully overlap, corresponding to the same physical location and indistinguishable neural activity.

Furthermore, the splitting effect can be studied through the lens of two different models introduced in Figure 4-A. First, the “Temporal Context Model” (TCM) (Howard and Kahana, 2002) emphasizes sensitivity to prior experiences, meaning the animal’s current neural state depends on its recent history. When recent pasts are similar, population activity at a given location exhibits closer proximity in PCA space. Conversely, when the recent past differs, the distances increase. This phenomenon is demonstrated by the varying distances between the markers C1, C2, and C3, each representing specific positions within the central corridor. When the animal reaches C1 in the maze, it has traversed the entire central corridor, resulting in a shared trajectory that accounts for the smaller distance between the C1 markers. In contrast, the animal’s past experiences at C3 differ significantly, which explains the greater distance between the C3 markers. Second, the “Latent State Inference” (LSI) (Gershman and Niv, 2010) organizes experiences into discrete states, resulting in a neural state less sensitive to historical context. As a result, the distances between the two neural trajectories in the central corridor remain constant: the distance between C1 markers is equal to the one between C2 markers and C3 markers. In the LSI model, experiences can be categorized into distinct states without strong dependencies on past experiences. For example, in scenarios where each stimulus is associated with a specific state, the categorization remains valid regardless of how previous stimuli were encountered. Duvelle et al. (2023) illustrates this with the example of one arm being rewarded while the other is not.

**Figure 4:**
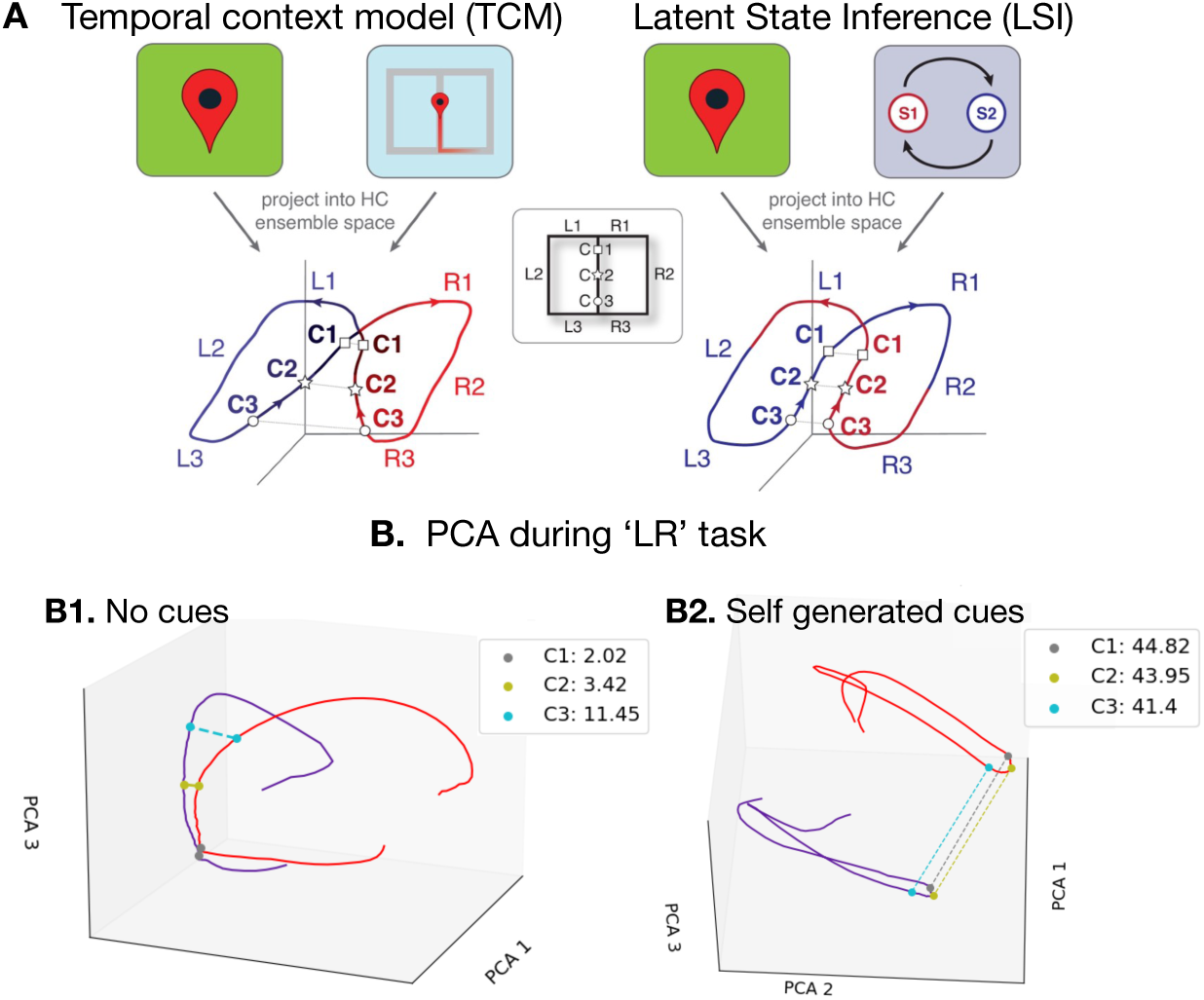
**A.** TCM vs. LSI. Figure from Duvelle et al. (2023). In the PCA space illustrating the TCM, the population activity at a specific location exhibits closer proximity in PCA space when the recent past is similar and greater distance when it is less similar. This is demonstrated in the distances between markers C1, C2, and C3. In the PCA illustrating the LSI, the distances between the markers C1, C2, and C3 don’t vary. The **B. PCA during the ‘LR’ task**. **B.1**. PCA applied to the regular model, resembling the TCM model. The increasing distances between C1, C2, and C3 markers suggest an amplified influence of past trajectories on neural activity. **B.2**. PCA applied to the self-generated cue model, resembling the LSI model. The distances between C1, C2, and C3 markers remain constant, indicating more distinct neural activity between the two trajectories due to the presence of the cues.

### Implicit approximation of the state-space environment and splitting effect at the population level

We proceed to a similar population analysis by applying 3D PCA to the reservoir states. This analysis was performed on all 10 seeds for both the regular and self-generated cue models. Examples from one seed of each model are shown in Figure 4-B1 and B2. The blue trace indicates the agent has just completed the left loop and is in the central corridor; the red trace corresponds to the agent having just completed the right loop. These two trajectories occupy distinct regions in the PCA space, despite they share the same location. The three markers (C1, C2, C3) indicating specific positions within the main corridor are distinctly separated with a measurable PCA distance, demonstrating the presence of splitter cells. Figures 5-A1 and B1 show the same PCA analysis while the bot traverses the central corridor across multiple loops. The PCAs show two linearly separable sub-attractors, corresponding to the two loops of the 8-shape. This result demonstrates the ESN’s ability to implicitly approximate the state-space of the 8-maze environment through continuous temporal dynamics. The model achieves this by mapping the input data into a higher-dimensional space (Jaeger, 2001). The recurrent connections within the reservoir enable the network to incorporate and reverberate past information, allowing for the encoding of temporal context and the integration of the past trajectory.

**Figure 5:**
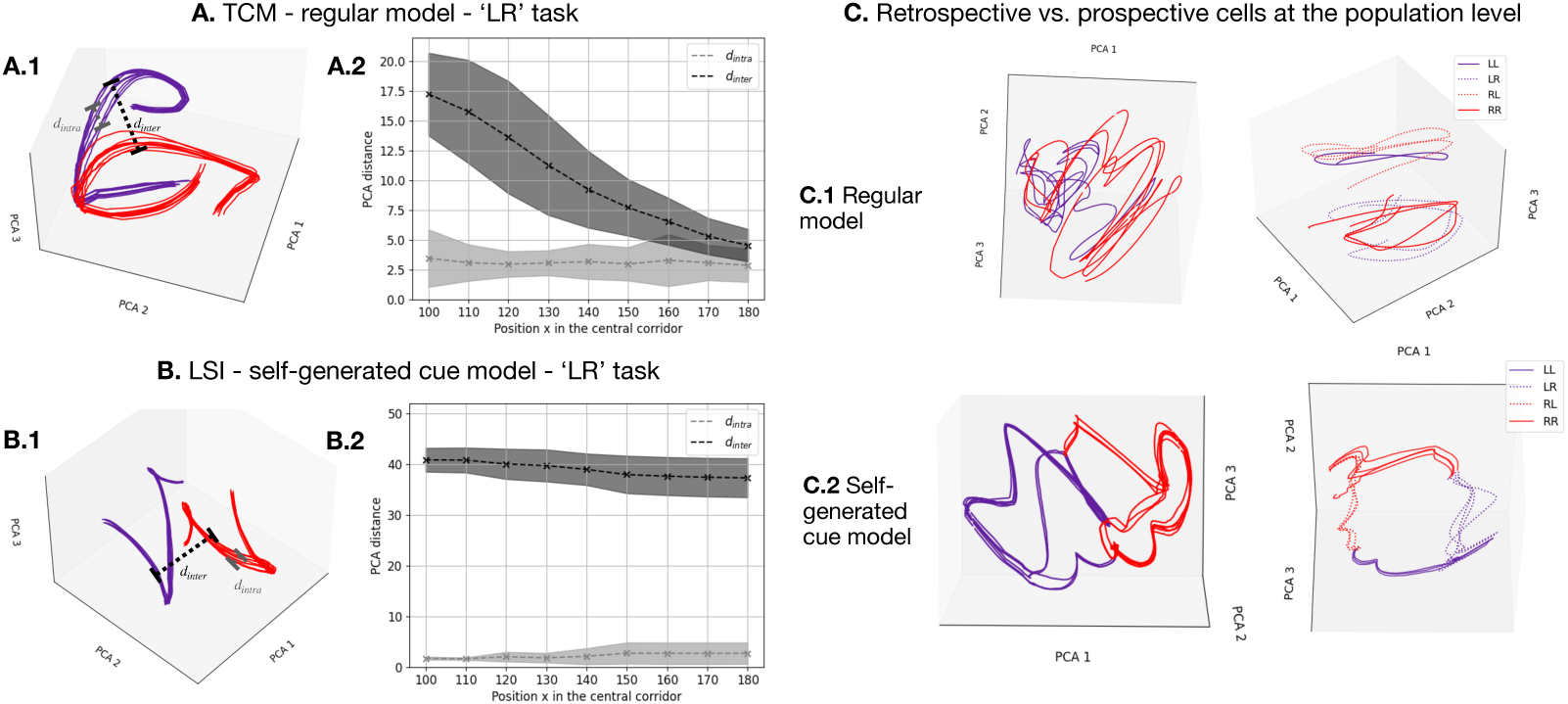
**A.** PCA during the ‘LR’ task over sevral trajectories. **A.1** Same as B.1 but for several loops. **A.2** PCA distance *d_intra_* between two identical trajectories (PCA distance between RL and RL, and LR and LR) and *d_inter_* between two different trajectories (PCA distance between RL and LR). The figure shows the average over 10 different seeds, each seed containing 15 passes per trajectory. We observe that *d_intra_ < d_inter_* until *d_inter_*becomes sufficiently small, consistent with the behavior expected under a TCM. This demonstrates that the TCM and LSI do not arise from variations in the bot’s lateral position, but rather because the two activity patterns originate from different trajectories **B.1** Same as B.2 but for several loops. **B.2** Same as C.2 but with the self-generated cue model. We observe that *d_intra_ < d_inter_* consistently across all positions. **C. PCA during the ‘LL-RR’ task. C.1** The regular model fails to distinguish LL from RL and RR from RL, consistent with its inability to complete the task. **C.2** The self-generated cue model clearly separates all four trajectories, aligning with its successful task performance.

### The regular model corresponds the TCM

In the regular model, we observed decreasing distances between C1,C2 and C3 markers in the central corridor (Figure 4-B1, Figure 5-A.1 and A.2), which aligns with the TCM (Figure 4-A). This is expected, as the model is influenced not only by current sensory inputs but also by recently experienced events, as the past trajectory of the bot is stored through the reverberation of recurrently connected neurons, allowing for a context-dependent response to present events.

### The self-generated cue model corresponds the LSI

In contrast, in the self-generated cue model, the distances between neural trajectories increase or remain stable as the bot enters the central corridor (Figure 4-B2, Figure 5-B.1 and B.2), reflecting a stronger association with one of the two discrete states (right or left). The presence of contextual cues provides direction information at each time step within the central corridor, positioning the model in a more discrete configuration. Consequently, the model does not require extensive memory of the past trajectory to successfully complete the navigation task.

Figures 5-A2 and B2 show *d*_intra_, the PCA distance between identical trajectories (e.g., RL vs. RL and LR vs. LR), and *d*_inter_, the distance between different trajectories (RL vs. LR). Distances are averaged over 10 seeds, each comprising 15 passes per trajectory. At the C3 marker for the regular model (Figure 4-A2), and consistently across all positions for the self-generated cue model (Figure 5-B2), *d*_inter_ is significantly greater than *d*_intra_. This indicates that the differences in neural activity are driven by distinct underlying trajectories, rather than minor lateral variations within each trajectory.

### Differentiation of retrospective and prospective cells at the population level

Figure 5-C shows a 3D PCA of internal states during the RR-LL task for the regular model, which failed after two loops. The left subpanel spans the full maze; the right focuses on the central corridor. No clear separation is observed between trajectories. In particular, the model fails to distinguish LL from RL and RR from RL. This likely reflects insufficient memory capacity to represent the four required sequences, contributing to task failure. In contrast, the self-generated cue model, which successfully completed the task beyond five loops, shows well-separated neural trajectories in the 3D PCA, as illustrated in Figure 5-C2. In the right subpanel, the four trajectories are clearly distinguished within the central corridor, and the full-maze PCA in the left subpanel also reveals a better separation between the two loops. The separation between RR and LR, and between RL and LL, reflects retrospective coding, neuronal activity differs depending on past trajectories. Conversely, the separation between LL and LR, and between RR and RL, indicates prospective coding, activity differs based on upcoming paths.

Together, these findings support and extend the hypotheses proposed by Duvelle et al. (2023) regarding the properties of splitter cells. We observe TCM-like activity in the regular model and LSI-like activity in the self-generated cue model. Furthermore, the ability to successfully complete the task may be linked to the presence of a population-level splitting effect. In cases where the model fails to generate distinct neural trajectories, it also fails to accomplish the task, reinforcing the importance of trajectory differentiation in successful performance. Notably, even when sensory cues reduce memory demands, splitter cells still appear, providing further support that they are likely a byproduct of the behavior itself and emerge as long as the task is performed, regardless of whether they are functionally required.

### Decoding position, decision, and head direction from the internal state of the reservoir

Three readouts were added to the reservoir to extract additional spatial information as the bot navigated, referred as decoder readouts. The first decoder accurately identified the bot’s position with 96% accuracy, consistently activating the correct location on the map (Figure 6-A). The second decoder achieved 94% accuracy in estimating head direction, confirming that orientation is also reliably embedded in the reservoir’s internal states (Figure 6-B). The third decoder predicted the bot’s next decision with 84% accuracy. As shown in Figure 6-C, after a right turn, the reservoir state evolved to favor a left turn, indicating that internal dynamics anticipate future choices. These results demonstrate that spatial information of the bot can be decoded from the reservoir thanks to its capacity of implicitly approximate the state-space of the environment.

**Figure 6:**
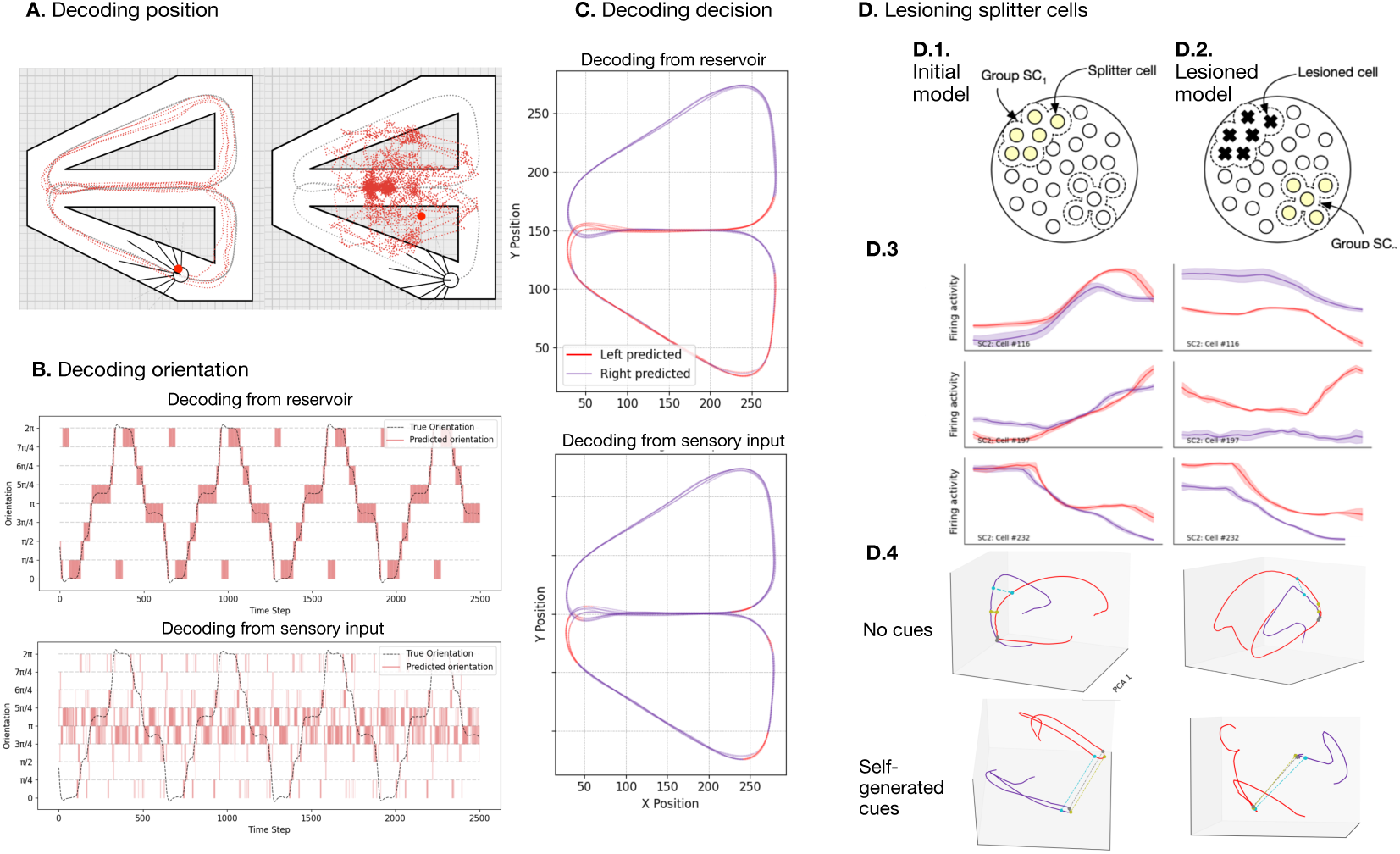
**A.** Snapshot of bot navigation and position predictions. The grey dotted line indicates the bot’s actual trajectory, while the red dotted line represents the predicted position at each time step based on the reservoir state. The decoder readout effectively tracks the bot’s position within the maze from the reservoir states (left) but not from the sensory input (right). **B. Orientation prediction.** The bot’s orientation is monitored over approximately 4000 time steps (black dotted line). The reservoir state enables highly accurate predictions of the bot’s next orientation (in red) from the resevoir states (top) but not from the sensory input (bottom). **C. Decision prediction**. Over 4000 time steps, the reservoir state reveals early internal shifts after the decision point. These shifts enable predictions within just a few dozen time steps, that the bot will switch direction at the next decision point (top). Predicting the next decision is not possible from the sensory inputs (bottom).**D. Silencing all splitter cells in the model**. **D.1-2**. Yellow cells represent splitter cells (SC), and those marked with a black cross are the silenced ones. First, all splitter cells are identified in the full model with or without cues (left). They are subsequently silenced, resulting in the lesioned model (right). The lesioned model is still able to perform the task, and a new set of splitter cells (SC2) emerges. **D.3. Activity of selected cells before and after lesioninf**. Each row shows the activity of the same neuron across both models. These cells did not exhibit a splitting effect before the lesion but developed one afterward. This change highlights how lesioning induces a reorganization of the model’s dynamics. **D.4. 3D PCA**.The splitting effect is still visible at the population level after lesion.

### Sensor-Based Decoding

In contrast, applying the same decoding techniques directly to raw sensor inputs failed to predict position, head-direction, or decision as depicted in Figure 4.-B,C and D. This confirms that the reservoir’s internal states encode critical space information, while the ambiguous sensory data alone does not.

### 3.5 Silencing splitter cells inside the models

#### Dynamic emergence of splitter cells supports persistent task performance

During the LR task, we identified an average of 217 splitter cells (SC1) in the regular model and 326 in the self-generated cue model across 10 seeds, as depicted in Table 3. All SC1 cells were subsequently silenced in both models, as illustrated in Figure 6-D.1 and D.2. The readout layers were re-trained using the same hyperparameters, without modifying the structural connectivity (see Figure 6-D.1). Despite SC1 inactivation, the lesioned models in both conditions successfully performed the alternation task across all seeds. On average, 256 new splitter cells (SC2) emerged in the lesioned regular model, and 216 SC2 in the lesioned self-generated cue model (see Table 3). Figure 6-D.3 shows examples of cells that were not initially splitter cells but became so after lesioning, and Figure 6-D.4 shows that the splitting effect at the population level is still observable after lesion. These findings indicate that silencing SC1 triggered a reorganization of the internal dynamics, giving rise to a new population of splitter cells (SC2). Thus, this result supports the idea that as long as the model performs the alternation task, splitter cells are likely to emerge from different neural populations. They reflect a flexible coding mechanism tied to task demands, rather than static circuit features. Similar re-organization was observed when silencing SC1 in the self-generated cue model during the LL-RR task, where 8 out of 10 seeds still succeeded.

**Table 3:** Average number of splitter cells across 10 seeds before and after lesioning the splitter cells during the ‘LR’ task.

| Model state | Regular Model | Self-Generated Cue Model |
| --- | --- | --- |
| Intact | $217.3 \pm 148.2$ | $326.7 \pm 138.5$ |
| Lesioned | $256.7 \pm 93.6$ | $216.2 \pm 64.8$ |

**Table 4:**
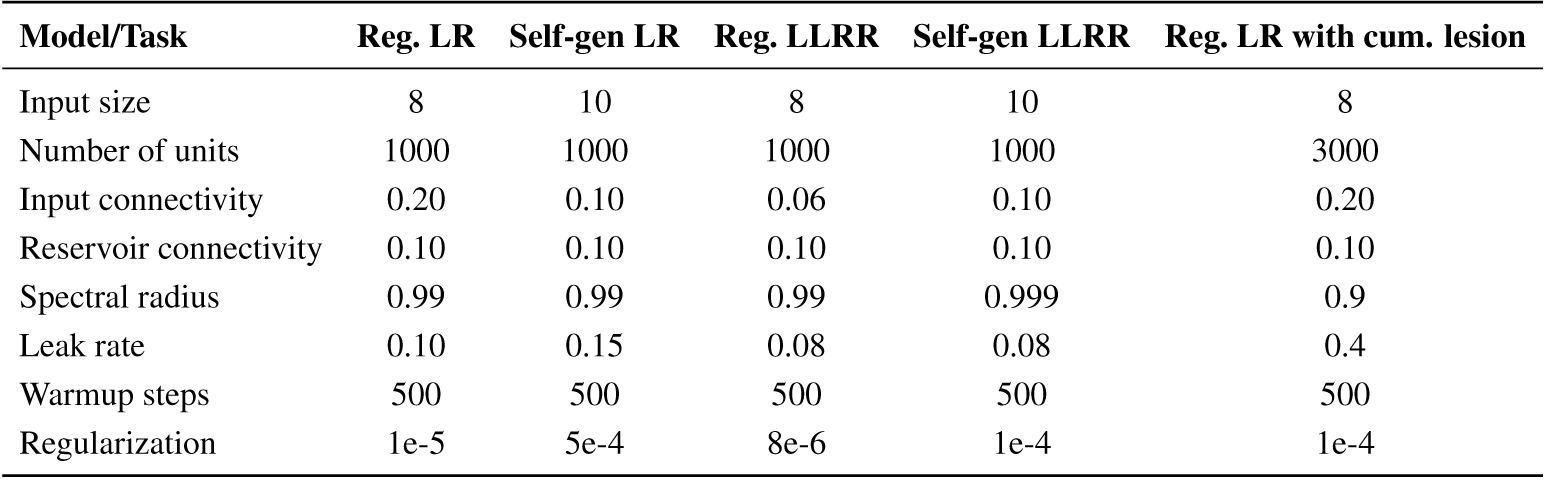
Model parameters when using offline linear regression learning.

#### Cumulative lesion of splitter cells

We conducted cumulative lesioning of splitter cells. Specifically, we repeatedly silenced all identified splitter cells, ran the model to allow new splitter cells to emerge, silenced them again, and repeated this process iteratively. To allow a sufficient number of lesion iterations before task failure, we used larger reservoirs of 3000 units and increased the leak rate to 0.4 to reduce the baseline number of splitter cells. The results, shown in Figure 7-A, begin with a reservoir of 3000 units, with each iteration reducing the reservoir size as splitter cells are cumulatively silenced. Remarkably,the agent successfully maintained task performance across up to nine successive lesion-retrain cycles: among the 30 random seeds tested, 3 failed to perform the task prior to any lesioning and were excluded; the remaining 27 seeds underwent sequential lesioning for a variable number of iterations, contingent on continued task performance following readout retraining after each lesion, as depicted in Figure 7-C. Notably, among the 131 successful reservoir states, 25 (19%) showed no detectable splitter cells following the lesion, while still maintaining successful navigation. This finding indicates that splitter cells are not strictly necessary for solving the trajectory-disambiguation task. To examine why some reservoirs solve the task without detectable splitter cells, we looked at the distribution of ANCOVA p-values across all reservoir units depicted in Figure 7-D. A reservoir with detected splitter cells shows a concentration of very low p-values, whereas a splitter-cell-free reservoir exhibits a more distributed p-value profile across units. This suggests that reservoirs without splitter cells may rather encode trajectory information through a distributed population of weakly selective units, none of which individually crosses the detection threshold of *p <* 0.0001.

**Figure 7:**
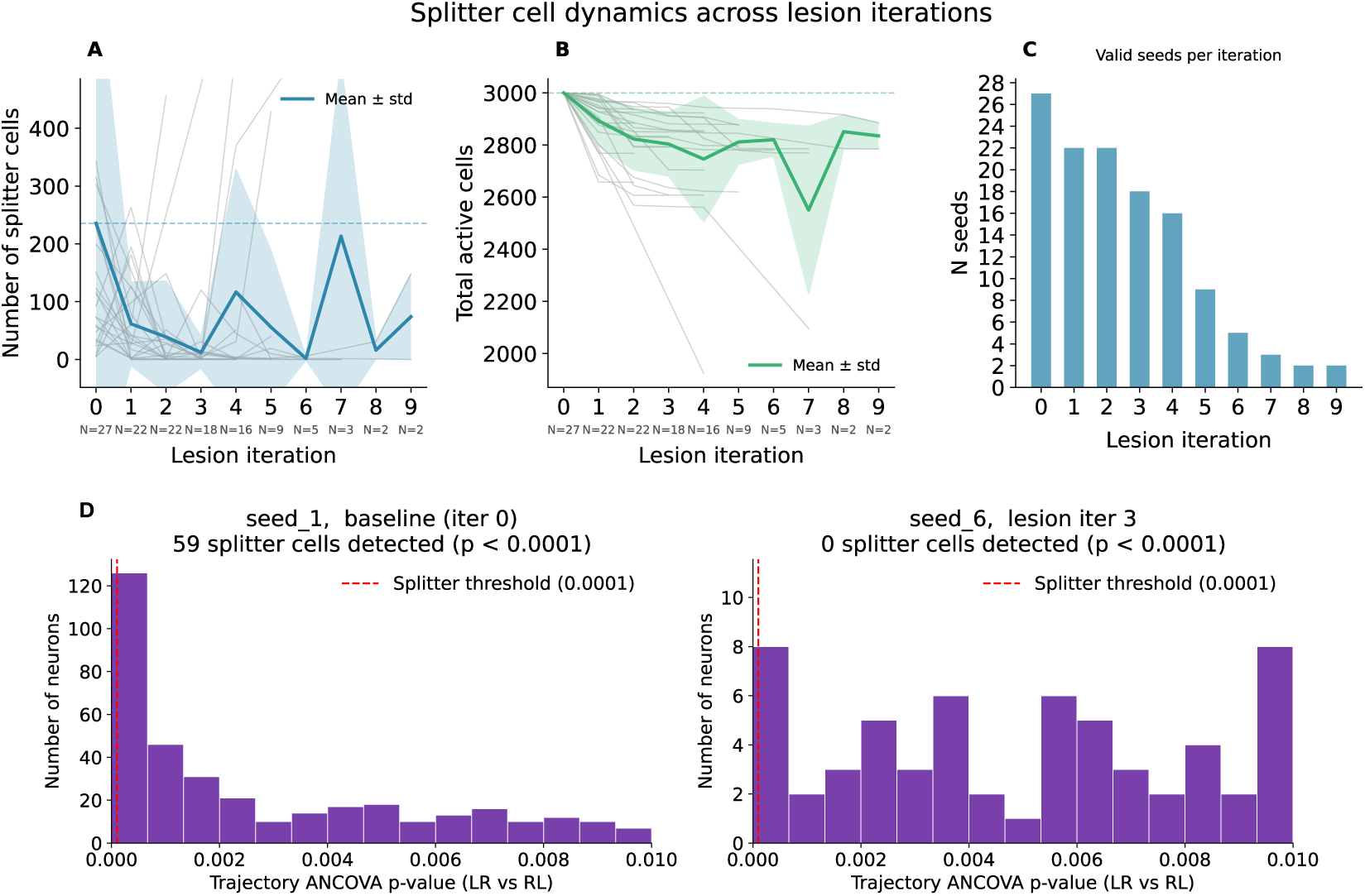
Cumulative lesion of splitter cells. Reservoirs were sequentially lesioned and retrained as long as they were able to successfully perform the R-L task for at least 15 loops. **A**. Number of splitter cells as a function of lesion iteration across 30 random seeds. **B**. Total number of active cells as a function of lesion iteration. The total number of active cells progressively decreased as lesions accumulated. **C**. Number of valid seeds at each lesion iteration. Of the 30 seeds tested, 27 were initially able to perform the task; 16 remained functional after 4 consecutive lesions, 8 after 5 cumulative lesions, and 2 after 9 lesions. **D**. Distribution of p-values across all neurons in two 3000-unit reservoirs, both capable of performing the task. Each bar represents the number of neurons whose ANCOVA test yielded a p-value within a given bin. Left: a reservoir in which 59 splitter cells are detected, showing a concentration of neurons with very low p-values below threshold. Right: a reservoir in which no splitter cells are detected by our criterion, yet p-values are continuously distributed across the full range rather than uniformly high, indicating the presence of weakly trajectory-selective neurons that individually fall below the detection threshold.

### Subspace alignment analysis across cumulative lesions

To further characterize the effect of cumulative lesioning on population geometry, we applied SVCCA (Raghu et al., 2017) to compare reservoir activity at baseline with each post-lesion iteration k. Figure 8-B summarizes the mean canonical correlation over the leading directions that explain 98% of the total correlation mass. This value remains high (0.8) across lesion iterations, indicating that the principal shared directions between intact and lesioned activity are largely preserved. Figure 8-D shows the full canonical correlation spectrum: *ρ_i_*1 for the first 7 modes, then declines, demonstrating that alignment is concentrated in a compact shared subspace rather than spread uniformly across all dimensions. Together, these results align with a global preservation of population geometry following lesions.

**Figure 8:**
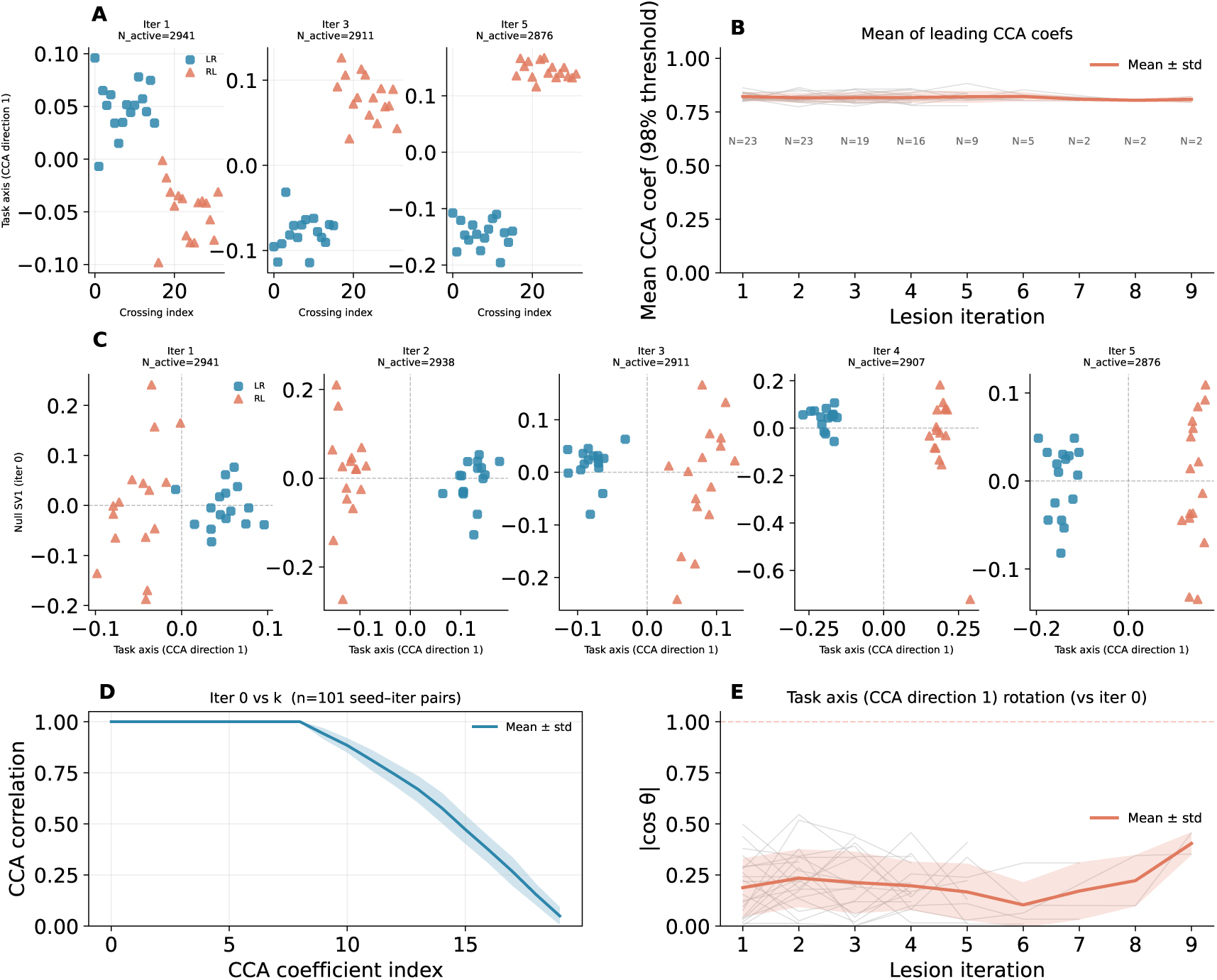
Subspace alignment after cumulative. **A**. Task axis projections for seed 1 across lesion iterations. LR and RL remain separated across iterations, indicating preserved task-related structure along the primary shared dimension between intact and lesioned networks. **B**. Mean canonical correlation over leading directions (98% cumulative *ρ* mass), comparing baseline vs lesion iteration k. Similarity stays around 0.8 across lesions, suggesting that dominant paired CCA directions are preserved. **C**. Task axis vs. baseline null activity across lesions. LR and RL remain separated on the task axis but overlap on null SV1, suggesting that this null dimension carries little task information. Variations along null SV1 across lesions indicate reorganization within the null subspace, while task structure along the potent axis is preserved. **D**. CCA correlation spectrum between iteration 0 and iteration k across all seeds and lesions. The x-axis shows the canonical direction index i (ordered by decreasing correlation), and the y-axis the canonical correlation *ρ_i_* between paired projections at iterations 0 and k. A plateau of 8 high-modes is followed by a sharp drop, suggesting that alignment is low-dimensional, with a few shared dimensions capturing most of the similarity. **E**. Task-axis rotation. *cos*(*θ*) is close to zero, indicating that the task axis rotates in neuron space and task information is re-encoded by a different neuronal combination.

Figure 8-A shows reservoir activity projected onto the task axis (CCA direction 1) for lesion iterations 1, 3, and 5 on an example seed. Each point represents one corridor crossing (LR or RL). LR and RL trajectories remain clearly separated across iterations, indicating that task-related structure is preserved on the primary dimension shared between intact and lesioned networks. Figure 8-C shows the same corridor crossings projected onto the current task axis (x-axis) and a fixed baseline null direction (y-axis), defined as the leading singular direction of iteration-0 activity after removing the mean task axis, held constant across all iterations. LR and RL overlap on the null direction, as expected for a baseline-defined null dimension, but the activity cloud shifts and spreads along this null direction across lesions, while separation on the task axis persists. This pattern is consistent with redistribution of activity within the null subspace rather than loss of task-relevant structure.

Figure 8-E shows that *cos*(*θ*) between the task axis at baseline and at iteration k re-mains approximately 0.2, corresponding to an angle of 78°, indicating that the task axis rotates substantially in neuron space across lesions. Task information is therefore reencoded by a different combination of neurons at each iteration, even though LR/RL separation and decoding accuracy remain preserved. Together, these results suggest that splitter re-emergence after lesion reflects a reorganization that preserves task-relevant population structure while redistributing activity in the null subspace, consistent with the theoretical framework of Churchland and Shenoy (2024), with the nuance that the task axis rotates in neuron space, meaning the same computation is implemented by a different neuronal combination at each iteration.

### Forced mode

Because the reservoir remains fixed during training and testing (fixed *W_in_* and *W*), training does not affect splitting activity. Instead of retraining for each condition, we forced the model to perform the task by providing the sensory inputs corresponding to correct alternation, enabling systematic evaluation across conditions and hyperparameters without optimization.

### The number of splitter cells depends on the reservoir hyperparameters

In forced mode, we tested different values of spectral radius, leak rate, and recurrent connectivity, examining their effects on both the number of splitter cells and decoder accuracy. Results are shown in Figure 9, based on simulations run over 10 seeds per hyperparameter setting, using the regular model on the LR task. Figure 9-A shows that the number of splitter cells increases with the spectral radius. Figure 9-B indicates an optimal leak rate around 0.15, corresponding to the peak number of splitter cells. It can be interpreted in terms of memory capacity. The leak rate controls the effective memory timescale of the reservoir: lower leak rates produce slower dynamics and longer memory, whereas higher leak rates produce faster but shorter-lived dynamics. The trajectory-disambiguation task requires the network to retain information about the past trajectory across the duration of central corridor traversal. A leak rate near 0.15 produces an optimal memory timescale allowing past-trajectory information to be retained long enough to influence neural activity within the central corridor. Leak rates much smaller or much larger than this optimum reduce the count of detectable splitter cells, through excessive memory persistence or insufficient memory respectively, but do not eliminate them entirely. In contrast, Figure 9-C demonstrates that the connectivity of *W* has no significant effect on splitter cell count. These trends align with decoder performance shown in Figure 9-D,E and F: position, decision, and orientation decoding are most accurate at the same leak rate that maximizes splitter cells, and decoding improves with increasing spectral radius. As with splitter cells, connectivity has no clear effect on decoding accuracy. Overall, these results suggest that specific reservoir properties, particularly spectral radius and leak rate critically influence the number of splitter cells and the quality of internal representations, while connectivity has little impact.

**Figure 9:**
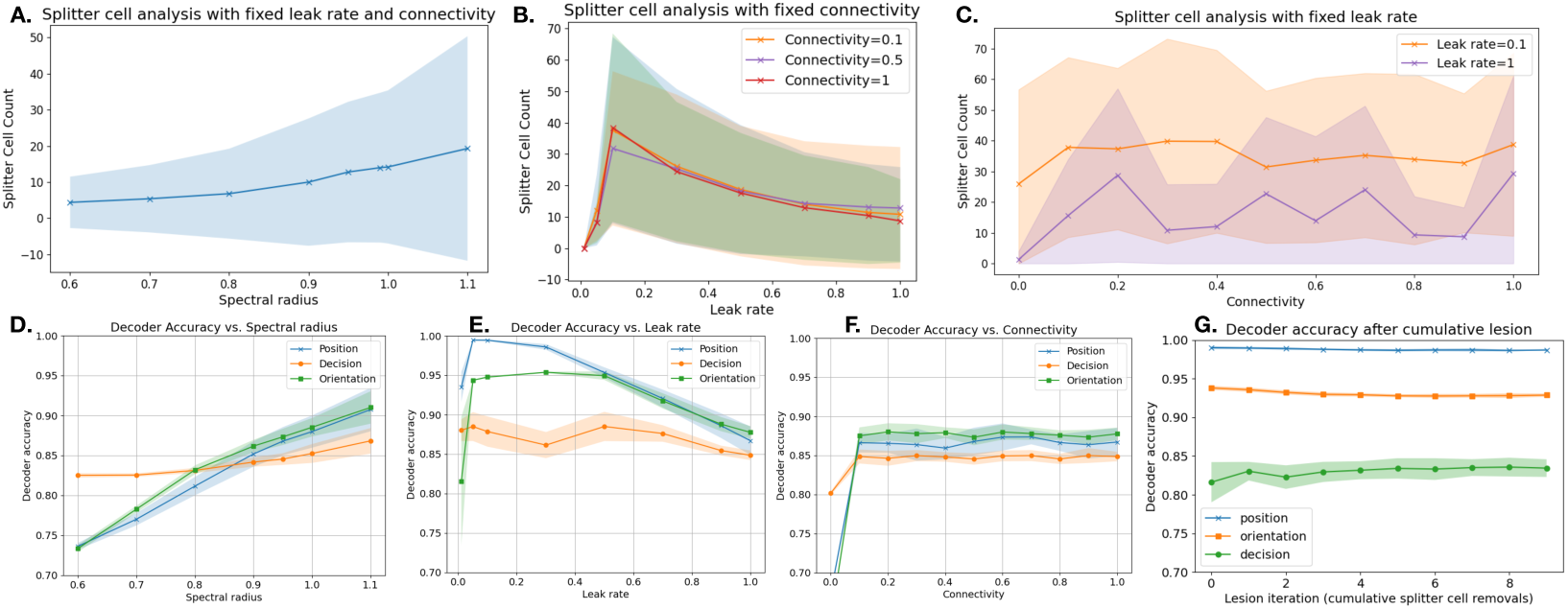
Evolution of splitter cell count and decoder accuracy across hyperparameters. Curves show the mean across 10 seeds, shaded areas represent standard deviation. Splitter cell count is shown as a function of **A**. spectral radius, **B.** leak rate, and **C**. reservoir connectivity. Decoder accuracy is shown as a function of **D**. spectral radius, **E**. leak rate, and **F**. reservoir connectivity.**G**. Decoder accuracy for position, decision and orientation following cumulative lesioning.

The consistent emergence of splitter cells support the idea that these representations are byproducts of correct task execution, rather than fixed prerequisites for behavior. This view aligns with Luo et al. (2024), who argue that cell types like place and grid cells may not be essential circuit components but instead arise inevitably from task constraints and behavior. In their study, spatial decoding also remained intact even after removing units classified as spatial cell types, indicating that these representations are neither unique nor necessary.

## 4 Discussion

The model under consideration is characterized by its fundamental simplicity, consisting of a random recurrent network that is informed by sensors enabling the computation of directional changes. Despite this apparent simplicity, the model is able to solve a continuous and non-trivial alternating navigation task, using ambiguous, limited and egocentric sensory information. Inside that model, we’ve been able to demonstrate for the existence of splitter cells Wood et al. (2000); Kinsky et al. (2020). By replicating their analyses on the reservoir, we identified activities indicative of splitter cells in rodents and demonstrated their re-emergence following targeted lesions and subsequent retraining. Furthermore, we’ve also validated the proposed theoretical framework by developing a model that incorporates concepts from both TCM (Temporal Context Mode), (Howard and Kahana, 2002) and LSI (Latent State Inference), (Gershman and Niv, 2010).

Several findings extend beyond this replication. First, splitter cells emerged even in configurations where working memory was not required for task performance. Second, across cumulative lesioning experiments, new splitter cells re-emerged in the vast majority of cases (81%). For these reservoirs, removing splitter cells without readout retraining produces immediate task failure: the existing readout had been calibrated to extract trajectory information via those specific neurons, and task-relevant information requires readout retraining to become accessible again, demonstrating informational redundancy. In the remaining 19% of reservoirs, the task was successfully solved without developing any detectable splitter cells, demonstrating that splitter cells are not a necessary computational solution in an absolute sense. Collectively, these findings imply that splitter cells are not functionally necessary for task execution but rather emerge as byproducts of it.

The existence of these splitter-cell-free reservoirs points to a deeper observation about the detection criterion itself. As noted by György Buzsáki (2019), neural response properties typically follow continuous distributions, and discrete cell-type labels inevitably reflect methodological thresholds imposed on that continuum. The ANCOVA criterion used here, a binary classification with p *<* 0.0001, is no exception: it applies a discrete label to a continuous distribution across the neuronal population. Reservoirs identified as splitter-cell-free by this criterion may nonetheless encode trajectory direction through a distributed population of weakly selective neurons, none of which crosses the detection threshold.

The concept of splitter cells bears a close resemblance to the notion of mixed selectivity (Rigotti et al., 2013), which refers to the property of neurons that respond to multiple, statistically independent task-relevant variables. Mixed selectivity is a well-established property of reservoir-type networks, emerging from random connectivity and nonlinear dynamics (Enel et al., 2016; Scleidorovich et al., 2022; Dominey, 2024), and is also observed in primate cortical neurons (Rigotti et al., 2010; Tye et al., 2024). Within this framework, splitter cells are units whose mixed selectivity weights trajectory-dependent variables strongly enough to be detected by our statistical criterion. Splitter cells re-emerge after lesion because surviving units carry partially overlapping selectivity profiles: removing the most strongly selective neurons does not eliminate the trajectory-relevant information from the population. Furthermore, some reservoirs solve the task without any detectable splitter cells because trajectory information is distributed across many weakly selective neurons, none of which individually crosses the detection threshold. The robustness of task performance across repeated lesion-retrain cycles is a direct consequence of the nature of mixed selectivity: the same task-relevant computation can be read out from multiple different subsets of the population.

Even though we did not claim any architectural plausibility with the hippocampal formation, the structure of the model is reminiscent of the CA3 structure with its highly recurrent nature, with the notable absence of learning inside the reservoir. In our case, the learning process is a simple linear identification of a random pattern of activity inside the reservoir in order to issue the proper motor command. This process is indeed well aligned with György Buzsáki (2019)’s inside-out view of the brain, where he suggests that brain activity is not generated from scratch during exploration and learning. According to this view, the brain can be considered as a vast reservoir capable of generating an extensive repertoire of neuronal patterns, initially independent of experience. In other words, these patterns have initially no meaning and acquire it through exploration and learning. In this sense, experience is primarily a process of matching preexisting neuronal dynamics to outside events, and learning does not create new brain activity from scratch, but rather selects preexisting pattern of activity responding to the external stimuli.

This “inside-out” view of brain function is supported by Langston et al. (2010), showing that spatial memory cells and their properties are already present when young pups first begin to move or even emerge early in postnatal development, before significant spatial exploration takes place (Muessig et al., 2016; Wills et al., 2014). These studies indicate that the hippocampus is capable of generating a wide range of possible neuronal trajectories even before the organism begins exploring its environment. On this regards, it is interesting to note that the randomness of the network in our model is not strongly constrained such that any matrix with the proper spectral radius will provide a sufficiently rich reservoir of dynamics from which the behavior can be acquired. Furthermore, in case of targeted lesion, it is relatively easy for the model to reacquire a behavior based on the new dynamics inside the reservoir, leading to the emergence of new splitter cells.

Beyond splitter cells, we also decoded the bot’s position, head direction, and upcoming decision from the activity units, information that echoes features of place cell, head direction cell, and decision cell activity. Even though we did not look for each and every other type of cells, we’re confident that we would probably be able to decode them. Remarkably, spatial and decision-related information remained decodable even after repeatedly lesioning all identified splitter cells in the model. This strongly suggests that their activities do not derive from structure nor learning (we have no learning, and no feedback inside the reservoir) but are rather indicative of a correlation with some pre-existing dynamics that are linked to the actual behavior. Considering for example decision cells (indicating decision to go left or right), it comes as no surprise that we can identify them inside the reservoir since the agent is really going left or right at the end of the central corridor, at least for an external observer. For the agent however, the sensory world is fully described by a set of eight sensors that do not convey the notions of *left* or *right*. Ultimately, this means that these decision cells may only exist in the eye of the observer. In other words, if a reservoir can solve a task with a minimal set of inputs, any variable that can be decoded from the activity of the reservoir derived from the resolution of the task itself as opposed to the specificity of structures involved in the resolution. This interpretation resonates with the CRISP model proposed by Cheng (2013), which argues that sequential activity patterns in CA3 are intrinsic to the network rather than imposed by external cortical inputs. According to this view, CA3 contains a reservoir of pre-established intrinsic sequences that serve as a substrate onto which external experiences are mapped via synaptic plasticity in feedforward projections, rather than through plasticity in recurrent CA3 connections.

Finally, during the past decade, the understanding of neural coding has gradually shifted from a population vector sum where a behavioral output is generated when the sum exceeds a threshold, to a more complex theory involving multidimensional neural activity space[4]. This theoretical framework distinguishes between an output-potent subspace, whose activity directly drives behavioral output, and a null space, whose activity does not directly affect output but is computationally essential. In this context, splitter cells reflect only a small partial dimension of the global encoding of the neuronal population. Our subspace alignment analysis directly addresses this framework: the task axis, the dimension separating LR and RL trajectories, is preserved across cumulative lesion iterations, while activity redistributes in the null subspace. Notably, the task axis rotates in neuron space across iterations, indicating that the same task-relevant computation is reencoded by a different neuronal combination after each lesion. This reorganization accounts for the emergence of new splitter cells: disturbing the multidimensional population space by silencing specific neurons induces a general reorganization of neural activity that affects the response of remaining neurons to the task, hence producing new detectable splitter cells. The preservation of task-relevant structure across this reorganization demonstrates that the output-potent subspace is maintained even as the null space redistributes, consistent with the flexibility-via-subspace hypothesis. If we demonstrated how this can be achieved using supervised offline and online training inside the model, the question of continuous and gradual adaptation remains open.

## 6 Code availability

The code associated with this work is publicly available at https://archive.softwareheritage.org/browse/origin/directory/? origin url=https://github.com/naomichx/splitter cells in reservoir 2025

